# Differential temporal dynamics during visual imagery and perception

**DOI:** 10.1101/226217

**Authors:** Nadine Dijkstra, Pim Mostert, Floris P. de Lange, Sander Bosch, Marcel A. J. van Gerven

**Affiliations:** Radboud University, Donders Institute for Brain, Cognition and Behaviour, Nijmegen, the Netherlands

## Abstract

Visual perception and imagery rely on similar representations in the visual cortex. During perception, visual activity is characterized by distinct processing stages, but the temporal dynamics underlying imagery remain unclear. Here, we investigated the dynamics of visual imagery in human participants using magnetoencephalography. We show that, contrary to perception, the onset of imagery is characterized by broad temporal generalization. Furthermore, there is consistent overlap between imagery and perceptual processing around 150 ms and from 300 ms after stimulus onset, presumably reflecting completion of the feedforward sweep and perceptual stabilization respectively. These results indicate that during imagery either the complete representation is activated at once and does not include low-level visual areas, or the order in which visual features are activated is less fixed and more flexible than during perception. These findings have important implications for our understanding of the neural mechanisms of visual imagery.

---

Visual imagery is the ability to generate visual experience in the absence of the associated sensory input. This ability plays an important role in various cognitive processes such as (working) memory, spatial navigation, mental rotation, and reasoning about future events^1^. When we engage in visual imagery, a large network covering parietal, frontal and occipital areas becomes active^2,3^. Multivariate fMRI studies have shown that imagery activates similar distributed representations in the visual cortex as perception for the same content^4–6^. There is a gradient in this representational overlap, in which higher, anterior visual areas show more overlap between imagery and perception than lower, posterior visual areas^5,7^. The overlap in low-level visual areas furthermore depends on the amount of visual detail required by the task^8,9^ and the experienced imagery vividness^6,10^. Thus, much is known about the spatial features of neural representations underlying imagery. However, the temporal features of these representations remain unclear.

In contrast, the temporal properties of perceptual processing are well studied. Perception is a highly dynamic process during which representations change rapidly over time before arriving at a stable percept. After signals from the retina reach the cortex, activation progresses up the visual hierarchy starting at primary, posterior visual areas and then spreading towards secondary, more anterior visual areas over time^11–13^. First, simple features such as orientation and spatial frequency are processed in posterior visual areas^14^ after which more complex features such as shape and eventually semantic category are processed more anteriorly^15–17^. After this initial feedforward sweep, feedback from anterior to posterior areas is believed to further sharpen the visual representation over time until a stable percept is achieved^18–20^.

However, the temporal dynamics of visual imagery remain unclear. During imagery, there is no bottom-up sensory input. Instead, visual areas are assumed to be activated by top-down connections from fronto-parietal areas^21,22^. Given the absence of bottom-up input, it is unlikely that activity begins at lower levels and then reaches higher levels later in time, like during perception. A more likely scenario is that during imagery, high-level representations in anterior visual areas are activated first, after which activity is propagated down to more low-level areas to fill in the visual details. This would be in line with the reverse hierarchy theory^23,24^. Alternatively, there may be no ordering such that during imagery, perceptual representations at different levels of the hierarchy are reinstated simultaneously.

In the current study, we investigated this question by tracking the neural representations of imagined and perceived stimuli over time. We combined magnetoencephalography (MEG) with multivariate decoding. First, we investigated the stability and recurrence of activation patterns over time using temporal generalization^25,26^. If different areas are activated after each other, activation patterns will be changing rapidly and thus will not generalize to other time points. If different levels are activated simultaneously, representations at the onset will already show high generalization to time points further away.

Furthermore, to be able to dissociate between a bottom-up and a top-down flow of activation through the visual hierarchy during imagery, we investigated at which time points during perception a classifier could generalize to imagery. If visual areas are activated in line with the visual hierarchy, earlier time points during perception should show overlap with earlier time points during imagery than later time points of perception. If instead there is a reversal in the direction, we would see the opposite: late time points during perception overlapping with early time points during imagery and vice versa.

## Results

### Behavioral results

Twenty-five participants executed a retro-cue task in which they perceived and imagined faces and houses and rated their experienced imagery vividness on each trial (see Fig. 1). Prior to scanning, participants filled in the Vividness of Visual Imagery Questionnaire, which is a measure of people’s imagery ability^27^. There was a significant correlation between VVIQ and averaged vividness ratings *(r* = −0.45, *p* = 0.02), which indicates that people with a higher imagery vividness as measured by the VVIQ also rated their imagery as more vivid on average during the experiment. Participants reported relatively high vividness on average (49.6 ± 26.6 on a scale between −150 and +150). There was no significant difference in vividness ratings between faces (54.0 ± 29.7) and houses (48.7 ± 26.7; *t*(24) = 1.46, *p* = 0.16). To ensure that participants were imagining the correct images, on 7% of the trials participants had to indicate which of four exemplars they imagined. The imagined exemplar was correctly identified in 89.8% (± 5.4%) of the catch trials, indicating that participants performed the task correctly. There was also no significant difference between the two stimulus categories in the percentage of correct catch trials (faces: 90.9 ± 6.6, houses: 88.8 ± 7.1; *t*(24) = −1.25, *p* = 0.22).

**Figure 1.**
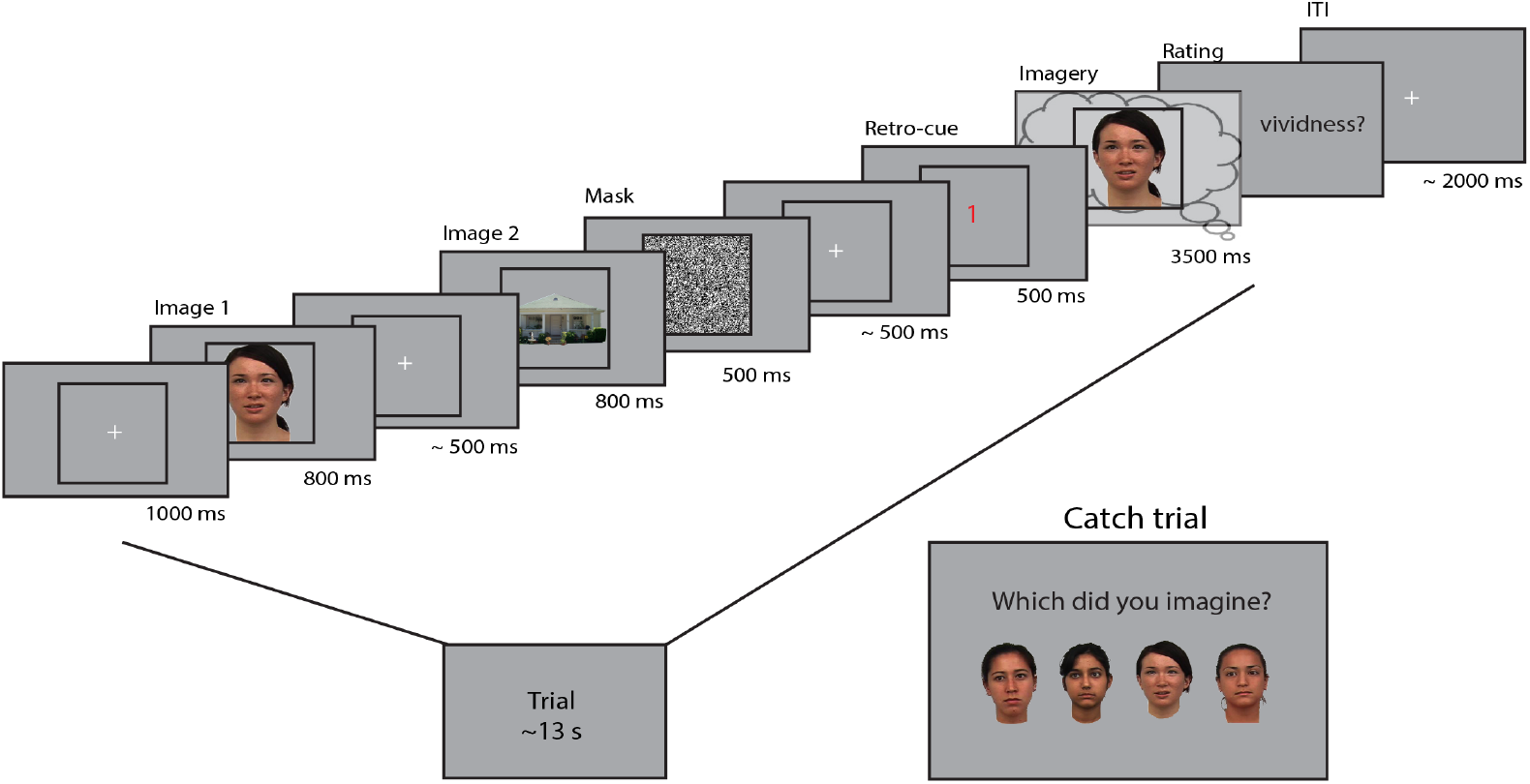
Experimental design. Two images were presented for 0.8 seconds each, with a random inter-stimulus interval (ISI) between 400 and 600 ms. After the second image, a mask with random noise was on screen for 500 ms. The retro-cue indicating which of the two images the participants had to imagine was shown for 500 ms. Subsequently, a frame was presented for 3.5 s within which the participants imagined the cued stimulus. After this, they rated their experienced vividness on a continuous scale. On a random subset (7%) of trials, the participants indicated which of four exemplars they imagined that trial.

### Representational dynamics during perception and imagery

To uncover the temporal dynamics of category representations during perception and imagery, we decoded the category from the MEG signal over time. The results are shown in Figure 2. Testing and training on the same time points revealed that during perception, significantly different patterns of activity for faces and houses were present from 73 ms after stimulus onset with the peak accuracy at 153 ms (Fig. 2A, left). During imagery, category information could be decoded significantly from 540 ms after retro-cue onset, with the peak at 1073 ms (Fig. 2A, right, Fig. S1B). The generation of a visual representation from a cue thus seems to take longer than the activation via bottom-up sensory input. Note that, to allow better comparison between perception and imagery, we only showed the first 1000 ms after cue onset during imagery (see Supplementary Figure S1 for the results throughout the entire imagery period).

**Figure 2.**
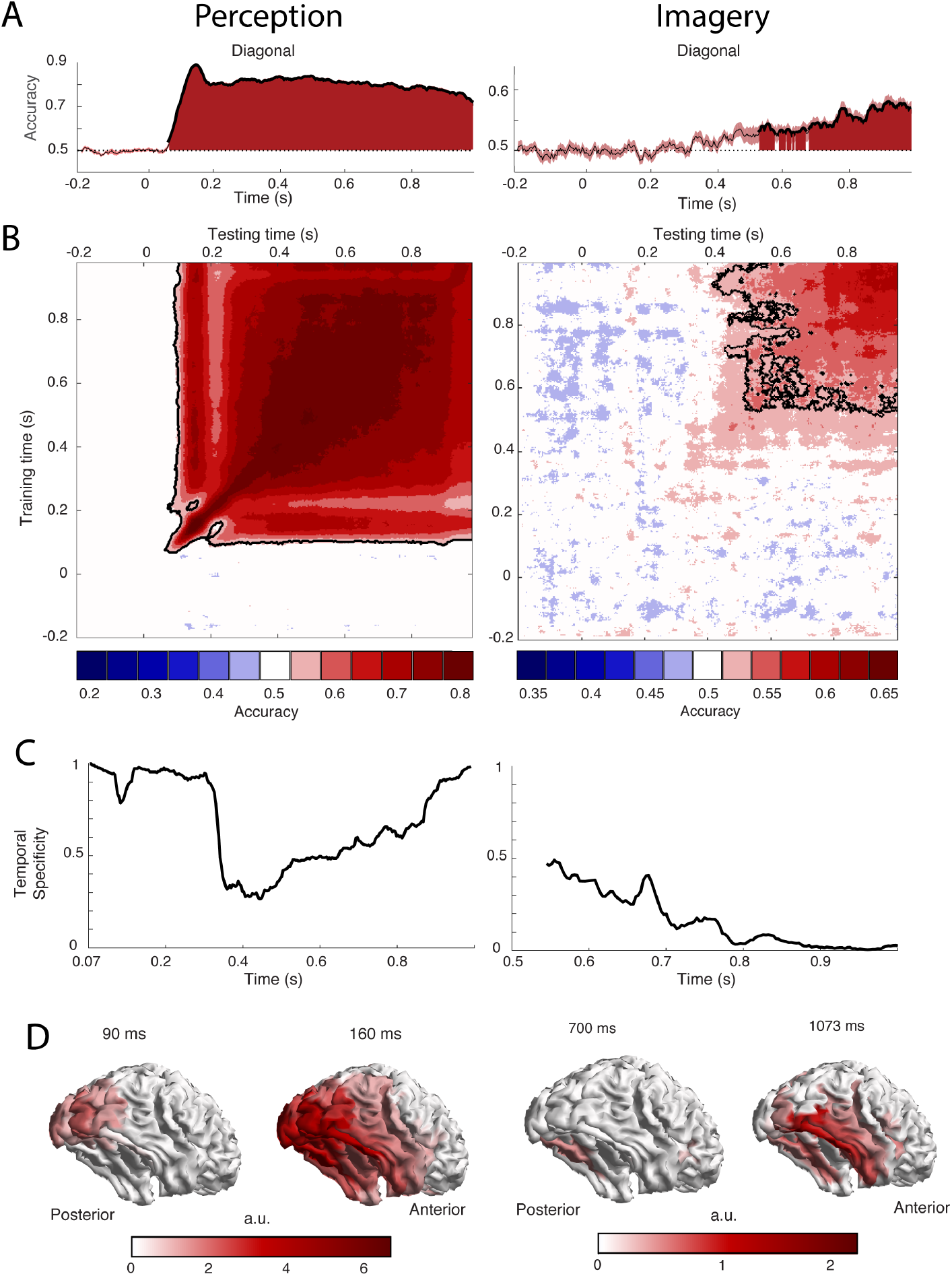
Decoding performance of perception and imagery over time. **(A)** Decoding accuracy from a classifier that was trained and tested on the same time points. Filled areas and thick lines indicate significant above chance decoding (cluster corrected, p < 0.05). The shaded area represents the standard error of the mean. The dotted line indicates chance level. For perception, zero signifies the onset of the stimulus, for imagery, zero signifies the onset of the retro-cue. **(B)** Temporal generalization matrix with discretized accuracy. Training time is shown on the vertical axis and testing time on the horizontal axis. Significant clusters are indicated by black contours. **(C)** Proportion of time points of the significant time window that had significantly lower accuracy than the diagonal, i.e. specificity of the neural representation at each time point during above chance diagonal decoding **(D)** Source level contribution to the classifiers at selected training times.

To reveal the generalization of representations over time, classifiers were trained on one time point and tested on all other time points [25] (Fig. 2B). Furthermore, to investigate the temporal specificity of the representations at each time point, we calculated the proportion of off-diagonal classifiers that had a significantly lower accuracy than the diagonal classifier of that time point^26^ (Fig. 2C; see *Materials and Methods).*

During perception, distinct processing stages can be distinguished (Fig. 2B-C, left). During the first stage, between 70 ms and 120 ms, diagonal decoding was significant and there was very high temporal specificity. This indicates sequential processing with rapidly changing representations^25^. During this time period, the classifier mostly relied on activity in posterior visual areas (Fig. 2D, left). Therefore, these results are consistent with initial feedforward stimulus processing. In the second stage, around 160 ms, the classifier generalized to neighboring points as well as testing points after 250 ms. The associated sources are spread out over the ventral visual stream (Fig. 2D, left), indicating that high-level representations are activated at this time. In the third stage, around 210 ms, we again observed high temporal specificity (Fig 2C, left) and a gap in generalization to 160 ms (Fig. 2B, left). This pattern could reflect feedback to low-level visual areas. Finally, from 300 ms onwards there is a broad off-diagonal generalization pattern that also generalizes to time points around 160 ms and an associated drop in temporal specificity (Fig. 2B-C, left). This broad off-diagonal pattern likely reflects stabilization of the visual representation.

In contrast, during imagery, we did not observe any clear distinct processing stages. Instead, there was a broad off-diagonal generalization throughout the entire imagery period (Fig. 2B, right; Fig. S1A). Already at the onset of imagery decoding, there was high generalization and low specificity (Fig. 2B-C, right). This indicates that the neural representation during imagery remains highly stable^25^. The only change seems to be in decoding strength, which first increases and then decreases over time (Fig. S1B), indicating that either representations at those times are weaker or that they are more variable over trials. The sources that contributed to classification were mostly located in the ventral visual stream and there was also some evidence for frontal and parietal contributions (Fig. 2D, right).

### Temporal overlap between perception and imagery

To investigate when perceptual processing generalizes to imagery, we trained a classifier on one data segment and tested it on the other segment. We first trained a classifier during perception and then used this classifier to decode the neural signal during imagery (Fig. 3A-B). Already around 350 ms after imagery cue onset, classifiers trained on perception data from 160 ms, 700 ms and 960 ms after stimulus onset could significantly decode the imagined stimulus category (Fig. 3A). This is earlier than classification within imagery, which started at 540 ms after cue onset (Fig 2A, right). Considering that we have more data in this analysis, this difference may reflect differences in signal-to-noise ratio (SNR) between the two analyses.

**Figure 3.**
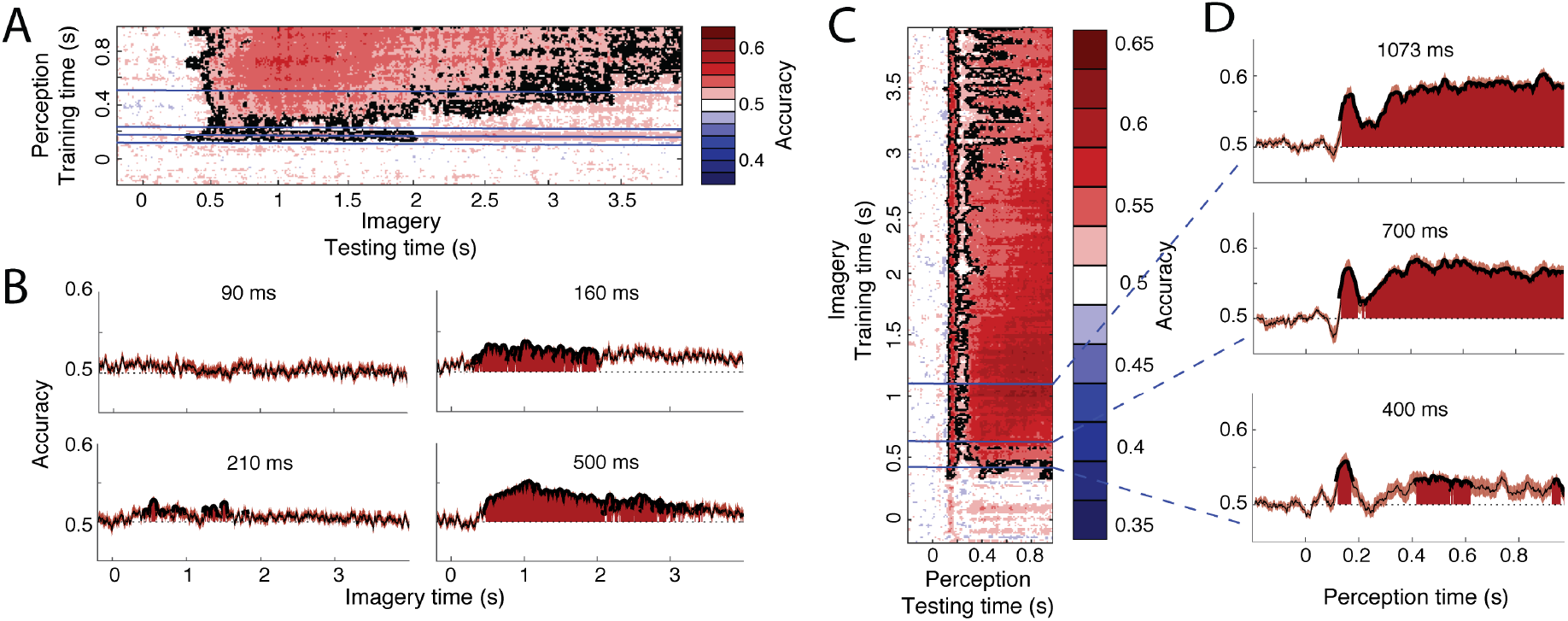
Generalization between perception and imagery over time. **(A)** Decoding accuracy from classifiers trained on perception and tested during imagery. The training time during perception is shown on the vertical axis and the testing time during imagery is shown on the horizontal axis. **(B)** Decoding accuracies for classifiers trained on the four stages during perception. **(C)** Decoding accuracy from classifiers trained on imagery and tested during perception. The training time during imagery is shown on the vertical axis and the testing time during perception is shown on the horizontal axis. **(D)** Decoding accuracies for different training times during imagery.

Furthermore, the distinct processing stages found during perception (Fig. 2B, left) were also reflected in the generalization to imagery (Fig 3A-B). Perceptual processes around 160 ms and after 300 ms significantly overlapped with imagery (Fig 3B, right plots). In contrast, processing at 90 ms did not generalize to any time point during imagery (Fig. 3B, top left). Perceptual processing at 210 ms showed intermittent generalization to imagery, with generalization at some time points and no generalization at other times (Fig. 3B, bottom left). Significant generalization at this time could also reflect the effects of smoothing over neighboring time points which are significant (see *Materials and Methods*). This would mean that there is no real overlap at 210 ms but that this overlap is caused by overlap from earlier or later time points.

To further pinpoint when perception started to overlap with imagery, we performed an additional analysis in which we reversed the generalization: we trained classifiers on different time points during imagery and used these to classify perception data. This analysis revealed a similar pattern of high overlap with perception around 160 and after 300 ms and low overlap before 100 ms and around 210 ms (Fig. 3C-D). Note that this profile is stable throughout imagery and is already present at the start of imagery, albeit with lower accuracies (Fig. 3-D, bottom panel). Furthermore, the onset of perceptual overlap is highly consistent over the course of imagery: overlap starts around 130 ms, with the first peak at approximately 160 ms (Fig. 3C). In general, cross-classification accuracy was higher when training on imagery than when training on perception (Fig. 3C vs. Fig. 3A). This is surprising, because training on high SNR data (in our case, perception) is reported to lead to higher classification accuracy than training on low SNR data^25^ (imagery). This difference may reflect the fact that the perceptual representation contained more unique features than the imagery representation, leading to a lower generalization performance when training on perception.

We also investigated whether the representational overlap between perception and imagery correlated with experienced imagery vividness. There were no significant correlations between classifier output and vividness ratings for any of the time points (see Supplementary Figure S2).

### Eye movements

Even though we attempted to remove eye movements from our data as well as possible (see *Materials and Methods*), it is theoretically possible that eye movements which systematically differed between the conditions caused part of the neural signal that was picked up by the decoding analyses^28^. In order to investigate this possibility, we tried to decode the stimulus category from the X and Y position of the eyes as measured with an eye tracker. The results for this analysis are shown in Supplementary Figure S3. During imagery, eye tracker decoding was at chance level for all time points, indicating that there were no condition-specific eye movements during imagery (Fig S3B). However, during perception, eye tracker decoding was significant from 316 ms onwards (Fig S3A), indicating that differences in eye movements between the conditions may have driven (part of) the brain decoding. If this were the case, there would be a high, positive correlation between eye tracker decoding and brain decoding. Figure S3C however shows that there was no such correlation, suggesting that our perception decoding results for that time window were not driven by eye movements.

## Discussion

We investigated the temporal dynamics of category representations during perception and imagery, as well as the overlap between the two. We first showed that whereas perception is characterized by high temporal specificity and distinct processing stages, imagery showed wide generalization and low temporal specificity from the onset. This indicates that the activity pattern picked up by the classifiers does not change much over the course of the imagery period. Furthermore, cross-decoding between perception and imagery revealed a very clear temporal overlap profile which was consistent throughout the imagery period. We found clear overlap between imagery and perceptual processing starting around 130 ms, decreasing around 210 ms and increasing again from 300 ms after stimulus onset. This pattern was already present at the onset of imagery.

These findings cannot be explained by a clear cascading of activity up or down the visual hierarchy during imagery. If there was a clear order in activation of different areas, we would not have observed such wide temporal generalization at the start of imagery but instead a more diagonal pattern, as during the start of perception^25^. Furthermore, we found that the complete overlap with perception is already present at the onset of imagery.

One interpretation of our results is that during imagery the complete stimulus representation, including different levels of the hierarchy, is activated simultaneously. However, there was no overlap between imagery and perceptual processing until 130 ms after stimulus onset, when the feedforward sweep is presumably completed and high-level categorical information is activated for the first time^29–31^. Overlap between perception and imagery in low-level visual cortex depends on the imagery task and experienced vividness ^5,6,8^. However, we did not observe a relationship between overlap at this time point and imagery vividness (Fig. S2). This absence of early overlap seems to imply that, even though early visual cortex has been implicated in visual imagery, there is no consistent overlap between imagery and early perceptual processing. One explanation for this discrepancy is that representations in low-level visual areas first have to be sharpened by feedback connections^31^ before they have a format that is accessible by top-down imagery. Alternatively, low-level visual activity during imagery may be more brief and variable over time than high-level activation, leading to a cancelling out when averaging over trials.

In line with this idea, an alternative explanation for our findings is that, in contrast to perception, the order in which visual features are activated during imagery is not fixed. This would be the case if, for example, participants sometimes first focused on the shape of the stimulus and then on the color, and sometimes first on the color and then the shape. High-level activations do show consistent activation during imagery, but perhaps the focus on specific visual features is more transient. Looking at time-locked processes will then cancel out these low-level activations, which does not happen when integrating over time with a method such as fMRI. This idea highlights the cognitive flexibility of imagery compared to perception. It would also explain why fMRI studies find that overlap between perception and imagery is decreased in low-level visual cortex compared to high-level visual areas^5^. In the current study, we cannot confidently dissociate the idea of a simultaneous onset of the complete visual representation or a changing spotlight on low-level features over time. More research into the temporal dynamics of imagery is needed to elucidate this issue.

The lack of generalization between imagery and perceptual processing around 210 ms after stimulus onset was unexpected. This time window also showed an increase in temporal specificity during perception, indicating rapidly changing representations. One possible interpretation is that around this time feedback from higher areas arrives in low-level visual cortex^32,33^. If low-level representations are indeed more transient, this would explain the decrease in consistent generalization. Another possibility is that processing at this time reflects an unstable combination of feedback and feedforward processes, which is resolved around 300 ms when representations become more generalized and again start to generalize to imagery. In line with this idea, processing from 300 ms after stimulus onset has been associated with percept stabilization^35–37^. Future studies looking at changes in effective connectivity over time are needed to dissociate these interpretations.

Surprisingly, we did not observe any influences of experienced imagery vividness on the overlap between perception and imagery over time (Fig. S2). One explanation for this is that we used whole-brain signals for decoding whereas the relationship between overlap and vividness has only been found for a specific set of brain regions^5,10^. Furthermore, the variability of low-level feature activations during imagery could prevent a consistent correlation between neural representations at specific times and imagery vividness. More studies on imagery vividness using MEG are necessary to explore this matter further.

In conclusion, our findings show that imagery is characterized by a different temporal profile than perception. Whereas perception showed high temporal specificity and distinct processing stages, imagery was characterized by a stable representational profile from the onset onwards. Furthermore, imagery showed consistent overlap with perceptual processing around 160 ms and from 300 ms onwards; times that are associated with the completion of the feedforward sweep and perceptual stabilization, respectively. This indicates that during imagery either the complete visual representation gets activated simultaneously and does not include low-level visual areas, or the order in which visual feature representations are activated is less fixed than during perception. It does not support the idea of a clear order of activation of the visual hierarchy during imagery, either bottom-up or top-down. Together, these findings reveal important new insights into the temporal dynamics of visual imagery and its relation to perception.

## Materials and methods

### Participants

Thirty human volunteers with normal or corrected-to-normal vision gave written informed consent and participated in the study. Five participants were excluded: two because of movement in the scanner (movement exceeded 15 mm), two due to incorrect execution of the task (less than 50% correct on the catch trials, as described below) and one due to technical problems. 25 participants (mean age 28.6, SD = 7.62) remained for the final analysis. The study was approved by the local ethics committee and conducted according to the corresponding ethical guidelines (CMO Arnhem-Nijmegen).

### Procedure and experimental design

Prior to scanning, participants were asked to fill in the Vividness of Visual Imagery Questionnaire (VVIQ): a 16-item questionnaire in which participants indicate their imagery vividness for a number of scenarios on a 5-point scale^27^. The VVIQ has been used in many imagery studies and is a well-validated measure of general imagery ability^5,6,10,38^. The score was summarized in a total between 16 and 80 (low score indicates high vividness). Subsequently, the participants practiced the experimental task for ten trials outside the scanner, after which they were given the opportunity to ask clarification questions about the task paradigm. If they had difficulty with the task, they could practice a second time with ten different trials.

The experimental task is depicted in Figure 1. We adapted a retro-cue paradigm in which the cue was orthogonalized with respect to the stimulus identity^39^. Participants were shown two images after each other, a face and a house, or a house and a face, followed by a retro-cue indicating which of the images had to be imagined. After the cue, a frame was shown in which the participants had to imagine the cued stimulus as vividly as possible. After this, they had to indicate their experienced imagery vividness by moving a bar on a continuous scale. The size of the scale together with the screen resolution led to discretized vividness values between −150 and +150. To prevent preparation of a motor response during imagery, which side (left or right) indicated high vividness, was pseudo-randomized over trials.

The face stimuli were adapted from the multiracial face database (courtesy of Michael J. Tarr, Center for the Neural Basis of Cognition and Department of Psychology, Carnegie Mellon University, http://www.tarrlab.org. Funding provided by NSF award 0339122). The house stimuli were adapted from the Pasedena houses database collected by Helle and Perona (California Institute of Technology). We chose faces and houses because these two categories elicit very different neural responses throughout the visual hierarchy, during both perception and imagery^3,40,41^, and are therefore expected to allow for high-fidelity tracking of their corresponding neural representations.

To ensure that participants were imagining the stimuli with great visual detail, both categories contained eight exemplars, and on 7% of the trials the participants had to indicate which of four exemplars they imagined (Fig. 1, Catch trial). The exemplars were chosen to be highly similar in terms of low-level features to minimize within-class variability and increase between-class classification performance. We instructed participants to focus on vividness and not on correctness of the stimulus, to motivate them to generate a mental image including all visual features of the stimulus. The stimuli encompassed 2.7 x 2.7 visual degrees. A fixation bull’s-eye with a diameter of 0.1 visual degree was on screen throughout the trial, except during the vividness rating. In total, there were 240 trials, 120 per category, divided in ten blocks of 24 trials, lasting about 5 minutes each. After every block, the participant had the possibility to take a break.

### MEG recording and preprocessing

Data were recorded at 1200 Hz using a 275-channel MEG system with axial gradiometers (VSM/CTF Systems, Coquitlam, BC, Canada). For technical reasons, data from five sensors (MRF66, MLC11, MLC32, MLF62, MLO33) were not recorded. Subjects were seated upright in a magnetically shielded room. Head position was measured using three coils: one in each ear and one on the nasion. Throughout the experiment head motion was monitored using a real-time head localizer^42^. If necessary, the experimenter instructed the participant back to the initial head position during the breaks. This way, head movement was kept below 8 mm in most participants. Furthermore, both horizontal and vertical electro-oculograms (EOGs), as well as an electrocardiogram (ECG) were recorded for subsequent offline removal of eye- and heart-related artifacts. Eye position and pupil size were also measured for control analyses using an Eye Link 1000 Eye tracker (SR Research).

Data were analyzed with MATLAB version R2017a and FieldTrip^43^ (RRID: SCR_004849). Per trial, three events were defined. The first event was defined as 200 ms prior to onset of the first image until 200 ms after the offset of the first image. The second event was defined similarly for the second image. Further analyses focused only on the first event, because the neural response to the second image is contaminated by the neural response to the first image. Finally, the third event was defined as 200 ms prior to the onset of the retrocue until 500 ms after the offset of the imagery frame. As a baseline correction, for each event, the activity during 300 ms from the onset of the initial fixation of that trial was averaged per channel and subtracted from the corresponding signals.

The data were down-sampled to 300 Hz to reduce memory and CPU load. Line noise at 50Hz was removed from the data using a DFT notch filter. To identify artifacts, the variance of each trial was calculated. Trials with high variance were visually inspected and removed if they contained excessive artifacts. After artifact rejection, on average 108 perception face trials (±11), 107 perception house trials (±12) and 105 imagery face trials (±16) and 106 imagery house trials (±13) remained for analysis. To remove eye movement and heart rate artifacts, independent components of the MEG data were calculated and correlated with the EOG and ECG signals. Components with high correlations were manually inspected before removal. The eye tracker data was cleaned separately by inspecting trials with high variance and removing them if they contained blinks or other excessive artifacts.

### Decoding analyses

To track the neural representations within perception and imagery, we decoded the stimulus category from the preprocessed MEG signals during the first stimulus and after the retro-cue for every time point. To improve the signal-to-noise ratio, prior to classification, the data were averaged over a window of 30 ms centered on the time point of interest. We used a linear discriminant analysis (LDA) classifier with the activity from the 270 MEG sensors as features (see ref. 44 for more details). A 5-fold cross-validation procedure was implemented where for each fold the classifier was trained on 80% of the trials and tested on the other 20%. To prevent a potential bias in the classifier, the number of trials per class was balanced per fold by randomly removing trials from the class with the most trials until the trial numbers were equal between the classes.

### Generalization across time and conditions

By training a classifier on one time point and then testing it on other time points, we were able to investigate the stability of neural representations over time. The resulting temporal generalization pattern gives information about the underlying processing dynamics. For instance, a mostly diagonal pattern reflects sequential processing of specific representations, whereas generalization from one time point towards another reflects recurrent or sustained activity of a particular process^25^. Here, we performed temporal generalization analyses during perception and during imagery to investigate the dynamics of the neural representations. Furthermore, to quantify the extent to which the representation at a given time point *t* was specific to that time point, we tested whether a classifier trained at time *t* and tested at time *t* (i.e. diagonal decoding) had a higher accuracy than a classifier trained at time *t*′ and tested at time *t* (i.e. generalization). This shows whether there is more information at time *t* than can be extracted by the decoder *t*′^26,45^. We subsequently calculated, for each time point, the proportion of time points that were significantly lower than the diagonal decoding, giving a measure of specificity for each time point. To avoid overestimating the specificity, we only considered the time window during which the diagonal classifiers were significantly above chance.

To investigate the overlap in neural representations between perception and imagery, a similar approach can be used. Here, we trained a classifier on different time points during perception and tested it on different time points during imagery and vice versa. This analysis shows when neural activity during perception contains information that can be used to dissociate mental representations during imagery and vice versa - i.e. which time points show representational overlap.

It has been shown that representational overlap between imagery and perception, as measured by fMRI, is related to experienced imagery vividness^5,6,10^. To investigate this in the current study, we related trial-by-trial vividness scores with classifier output over time as in ref. 46. An LDA classifier calculates for every trial the distance from a decision boundary that optimally discriminates the two classes. For classification, distances are subsequently transformed to a predicted class using a cut-off. This prediction is then used to calculate the percentage of correctly classified trials, i.e. the decoding accuracy^44^. However, the distance measure can be interpreted as the amount of evidence for a certain class, such that trials with more evidence for the correct class can be seen as having less ambiguous representations^46,47^. In the current context, the distance that is calculated by a classifier trained on perception can be interpreted as the degree of representational overlap with perception for that trial. Correlating these distances during imagery with experienced imagery vividness per time point reveals if, and at which points in time, overlap between perception and imagery relates to experienced vividness.

### Statistical testing

Decoding accuracy was tested against chance using two-tailed cluster-based permutation testing with 1000 permutations^48^. In the first step of each permutation, clusters were defined by adjacent points that crossed a threshold of p < 0.05. The t-values were summed within each cluster, but separately for positive and negative clusters, and the largest of these were included in the permutation distributions. A cluster in the true data was considered significant if its p-value was less than 0.05 based on the permutations. Correlations with vividness were tested against zero on the group level using the same procedure.

### Source localization

In order to identify the brain areas that were involved in making the dissociation between faces and houses during perception and imagery, we performed source reconstruction. In the case of LDA classifiers, the spatial pattern that underlies the classification reduces to the difference in magnetic fields between the two conditions (see ref. 49). Therefore, to infer the contributing brain areas, we performed source analysis on the difference ERF between the two conditions.

For this purpose, T1-weighted structural MRI images were acquired using a Siemens 3T whole body scanner. Vitamin E markers in both ears indicated the locations of the head coils during the MEG measurements. The location of the fiducial at the nasion was estimated based on the anatomy of the ridge of the nose. The volume conduction model was created based on a single shell model of the inner surface of the skull. The source model was based on a reconstruction of the cortical surface created for each participant using FreeSurfer’s anatomical volumetric processing pipeline (RRID: SCR_001847). MNE-suite (Version 2.7.0; RRID: SCR_005972) was subsequently used to infer the subject-specific source locations from the surface reconstruction. The resulting head model and source locations were co-registered to the MEG sensors.

The lead fields were rank reduced for each grid point by removing the sensitivity to the direction perpendicular to the surface of the volume conduction model. Source activity was obtained by estimating linearly constrained minimum variance (LCMV) spatial filters^50^. The data covariance was calculated over the interval of 50 ms to 1 s after stimulus onset for perception and over the entire segment for imagery. The data covariance was subsequently regularized using shrinkage with a regularization parameter of 0.01 (as described in ref. 52). These filters were then applied to the axial gradiometer data, resulting in an estimated two-dimensional dipole moment for each grid point over time. For imagery, the data were low-pass filtered at 30 Hz prior to source analysis to increase signal to noise ratio.

To facilitate interpretation and visualization, we reduced the two-dimensional dipole moments to a scalar value by taking the norm of the vector. This value reflects the degree to which a given source location contributes to activity measured at the sensor level. However, the norm is always a positive value and will therefore, due to noise, suffer from a positivity bias. To counter this bias, a permutation procedure was employed in which the noise was estimated and subtracted from the true data. Afterwards, the data were also divided by the noise estimates in order to counter the depth bias (for full details, see ref 52).

For each subject, the surface-based source points were divided into 74 atlas regions as extracted by FreeSurfer on the basis of the subject-specific anatomy^53^. To enable group-level estimates, the activation per atlas region was averaged over grid points for each participant. Group-level activations were then calculated by averaging the activity over participants per atlas region^54^.

## Additional information

### Acknowledgements

The authors would like to thank Marius Peelen for his comments on an earlier version of the manuscript and Jean-Rémi King for his suggestions for analyses.

## Supplementary material

**Figure S1.**
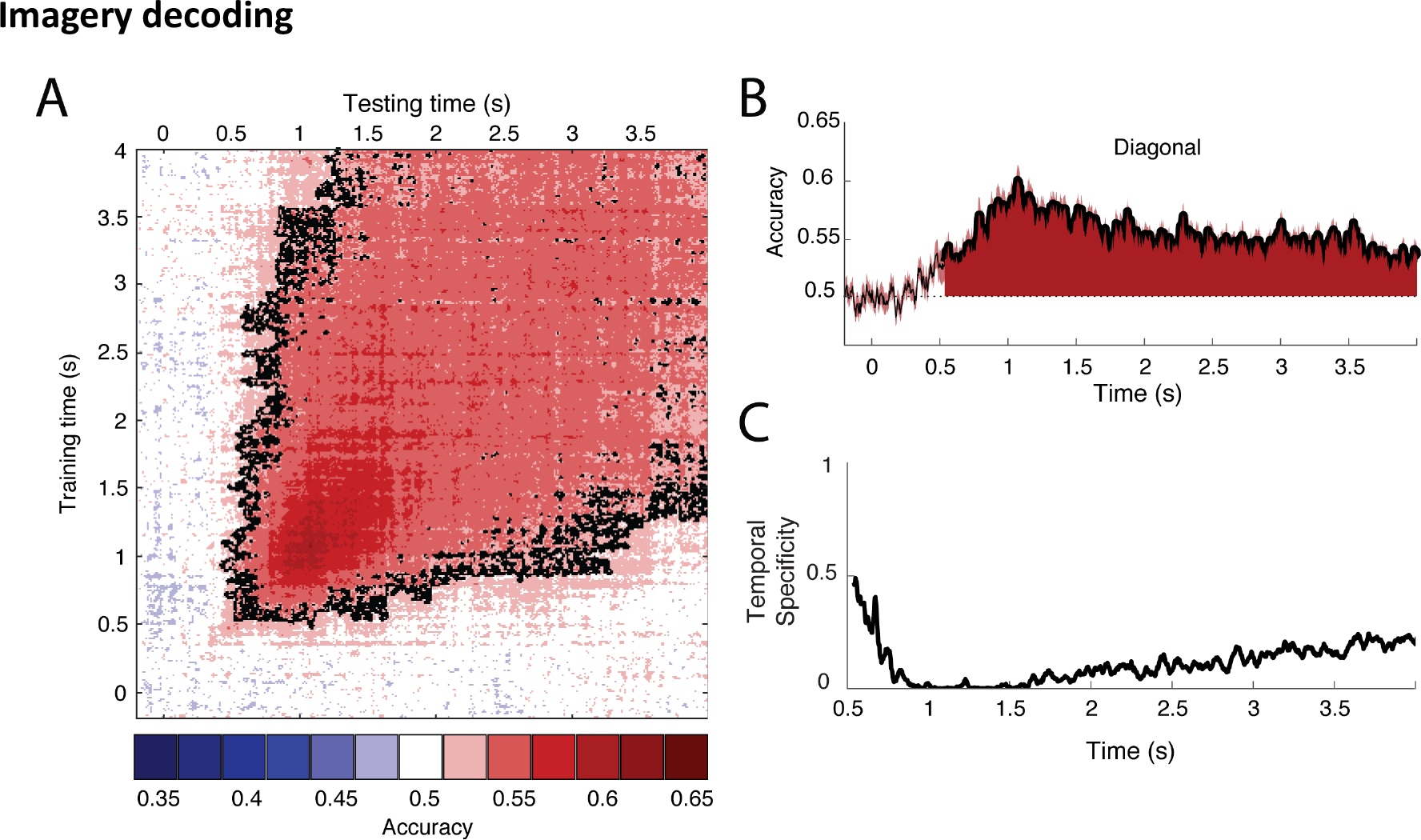
Decoding results throughout the entire imagery period. **(A)** Temporal generalization matrix. Training time is shown on the vertical axis and testing time on the horizontal axis. **(B)** Decoding accuracy from a classifier that was trained and tested on the same time points during imagery. **(C)** For each testing point, the proportion of time points that resulted in significantly lower accuracy than the diagonal decoding at that time point, i.e. the temporal specificity of the representations over time.

**Figure S2.**
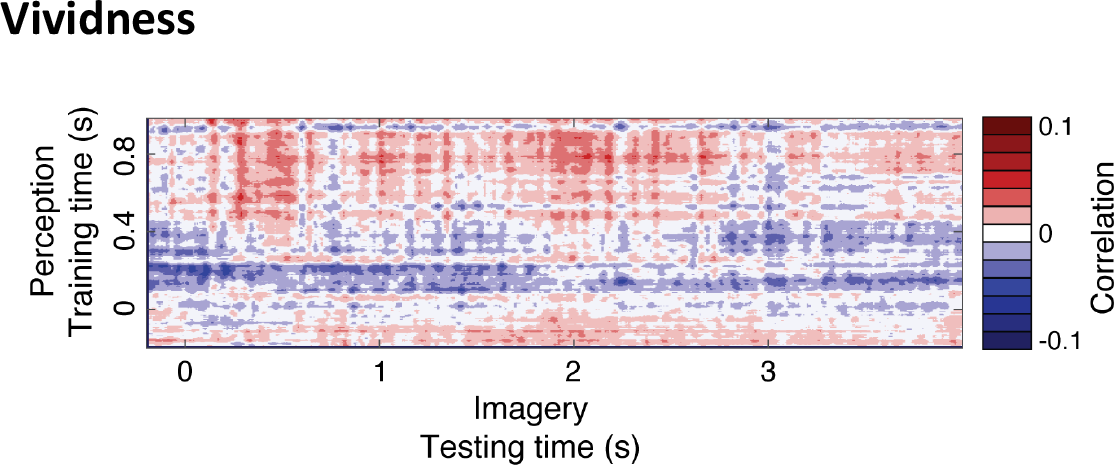
Correlation between vividness and classifier predictions. The calculated distances per trial during imagery of classifiers trained on perception were correlated with vividness ratings for every time point. Group averaged correlations were subsequently tested against zero using cluster based permutation testing. No significant clusters were found.

**Figure S3.**
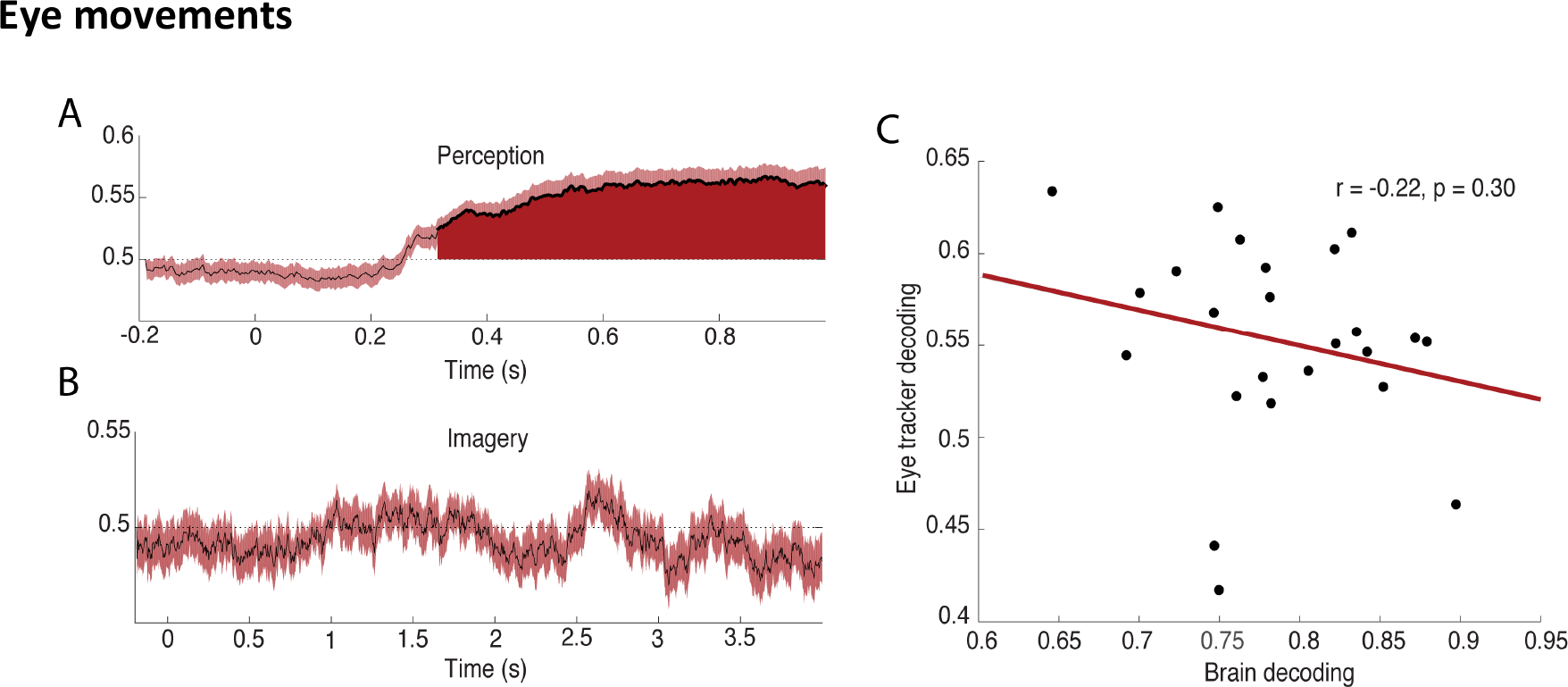
Decoding on eye tracker data. **(A)** Decoding accuracy over time on eye tracker data during perception. Filled areas and thick lines indicate significant above chance decoding (cluster corrected, p < 0.05). The shaded area represents the standard error of the mean. The dotted line indicates chance level. **(B)** Decoding accuracy over time on eye tracker data during imagery. **(C)** Correlation over participants between eye tracker decoding accuracy and brain decoding accuracy, averaged over the period during which eye tracker decoding was significant.

